# Teichoic acids anchor distinct cell wall lamellae in an apically growing bacterium

**DOI:** 10.1101/714758

**Authors:** Eveline Ultee, Lizah T. van der Aart, Dino van Dissel, Christoph A. Diebolder, Gilles P. van Wezel, Dennis Claessen, Ariane Briegel

## Abstract

The bacterial cell wall is a dynamic, multicomponent structure that provides structural support for cell shape and physical protection from the environment. In monoderm species, the thick cell wall is made up predominantly of peptidoglycan, teichoic acids and a variety of capsular glycans. Filamentous monoderm Actinobacteria, such as *Streptomyces coelicolor*, incorporate new cell wall material at the apex of their hyphal cells during growth. In this study we use cryo-electron tomography to reveal the structural architecture of the cell wall of this bacterium. Our data shows a density difference between the apex and subapical regions of chemically isolated sacculi. Removal of the teichoic acids with hydrofluoric acid reveals a rough and patchy cell wall and distinct lamellae in a number of sacculi. Absence of the extracellular glycans poly-β-1,6-*𝒩*-acetylglucosamine and a cellulose-like polymer, produced by the MatAB and CslA proteins respectively, results in a thinner sacculus and absence of lamellae and patches. Extracellular glycans might thus form or lead to the formation of the outer cell wall lamella. Based on these findings we propose a revisited model for the complex cell wall architecture of an apically growing bacterium, in which the network of peptidoglycan together with extracellular polymers is structurally supported by teichoic acids.

## Introduction

Bacteria are successful organisms that thrive in most environments. They withstand challenging conditions by synthesizing a stress-bearing cell wall, which provides rigidity to cells and defines their shape. Since the cell wall is in most cases essential for the cells’ survival, it is a prime target for antibiotic treatment. The core component of the cell wall is formed by a mesh of N - acetylglucosamine (GlcNAc) and N-acetylmuramic acid (MurNAc) glycan chains, which are cross-linked by peptide stems to form the peptidoglycan (PG) sacculus ^1^. Bacteria have been classified into two groups based on their cell envelope architecture: monoderm bacteria have a cell envelope that consists of a cytoplasmic membrane and a multilayered envelope, consisting of PG, teichoic acids (TAs) ^2,3^ and a variety of capsular glycans ^4–7^. Teichoic acids are long, anionic polymers and can be classified into wall teichoic acids (WTAs) and lipoteichoic acids (LTAs), based on their linkages to either the PG or lipid membrane respectively ^8,9^. They contain similar repetitive units linked by phosphodiester linkages which make up the long structure, and render the cell envelope negatively charged ^9,10^. In contrast, diderm bacteria have a thin PG layer that this is positioned between a cytoplasmic membrane and an additional outer membrane containing lipopolysaccharides (LPS) ^11^.

In order for a bacterial cell to grow, the PG needs to expand and be remodeled by the insertion of newly synthesized PG strands into the pre-existing sacculus. The cell wall can expand either by insertion of new cell wall material by lateral expansion, or by tip growth (Figure 1A). In many unicellular rod-shaped bacteria, such as in the model organisms *Escherichia coli* and *Bacillus subtilis*, the cell wall expands by the incorporation of new PG along the length of the cell ^12,13^. This lateral elongation is guided by highly curved MreB filaments along the cell circumference ^14–17^. In contrast, other bacteria, including actinobacteria and *Agrobacterium* species, grow by polar cell wall elongation, which is independent of MreB ^18,19^. In actinobacteria, the curvature sensitive protein DivIVA localizes to the cell poles and recruits the cell wall synthesis machinery ^20–22^. The current model of tip growth involves *de novo* PG synthesis at the apex guided by DivIVA and modification of the new wall material by L-,D-transpeptidases along the lateral wall ^23^.

**Figure 1.**
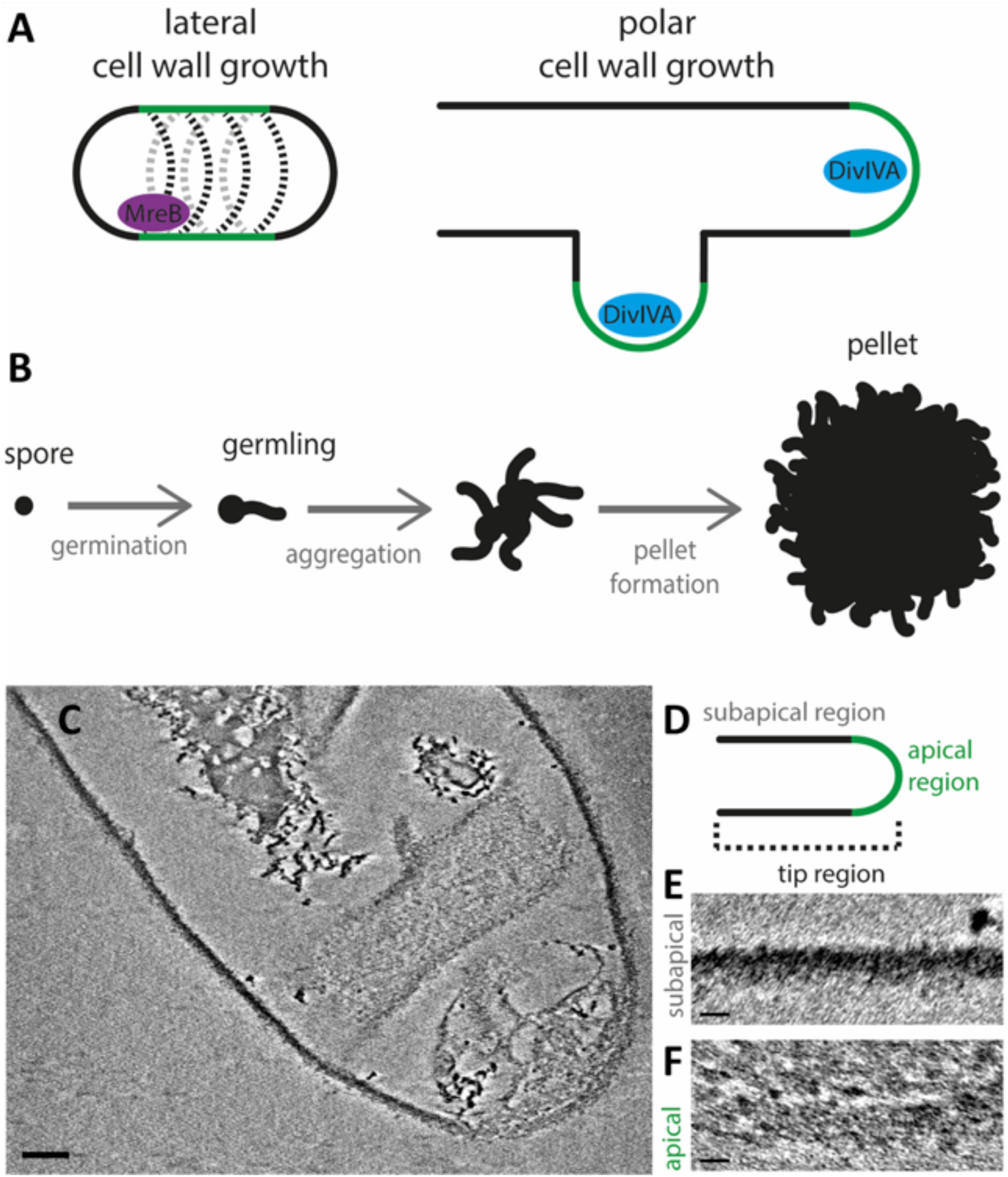
Model of *Streptomyces* growth and cryo-ETs of isolated *S. coelicolor* sacculi. The cell wall of bacteria is incorporated either at the lateral sides via the MreB filaments or at the apex driven by DivIVA. These processes are termed lateral and polar cell wall growth respectively (A). Panel B shows the growth and development of *Streptomyces coelicolor* spores in liquid medium, where the spores germinate, aggregate and form dense pellets facilitated by an extracellular matrix (B). Panel C shows an cryo-electron tomography slice of the tip region of a *S. coelicolor* sacculus with insets displaying differences in cell wall structure in the apical (E) and subapical regions (F), as depicted in the schematic (D). Scale bar (A) 100 nm, scale bars insets (E,F) 20 nm.

To address the organization of the cell envelope architecture of polar growing bacteria we studied the model organism *Streptomyces coelicolor* ^24^. *Streptomyces* species are soil-dwelling and multicellular actinobacteria that form long, branched hyphal cells, which collectively form an extended mycelial network ^25,26^. When grown in liquid, the hyphae form dense mycelial pellets by producing a glue-like extracellular matrix (Figure 1B) ^27^. This extracellular matrix is composed of a large variety of polymers ^4,28,29^. Two of these polymers have been characterized as key elements for pellet formation : on eispoly - β - 1, 6 - *𝒩* - acetylglucosamine (PNAG), produced by the MatAB proteins ^30,31^. The other one is a cellulose-like glycan, formed by the cellulose synthase-like enzyme CslA, which operates in conjunction with GlxA and DtpA at the hyphal tips ^32–35^. They were shown to add to the stability of the tip and are important for invasive growth ^32,36^. Deletion of either the *matAB* cluster (SCO2963 and SCO2962) ^30,31^ or *cslA* (SCO2836) ^32^ results in an open-growth morphology, which is characterized by the absence of pellet formation in liquid-grown media.

As the tips of growing hyphae continuously incorporate nascent PG, the architecture of the sacculus is likely structurally distinct from the mature cell wall at the lateral sides of the hyphae. Here we reveal structural differences of the cell wall at both the lateral sides and the nascent cell wall at the hyphal tip by applying cryo-electron tomography (cryo-ET) using a Volta Phase Plate (VPP) to allow high resolution imaging with high contrast.

The results of this study provide unprecedented insight into the architecture of the cell wall of a polar growing bacterium. The data suggest that the cell wall of *S. coelicolor* is comprised of two distinct lamellae. The inner lamella is composed of PG, and the glycans produced by the CslA and MatAB proteins form a discrete outer lamella, which is tethered to the inner layer via teichoic acids. Collectively, these findings warrant a revised model for the cell wall architecture in a polar growing bacterium.

## Results

### Isolated *S. coelicolor* sacculi reveal density differences in growing hyphal tip regions

In order to reduce the thickness of the sample and achieve high resolution cryo-electron tomograms of the cell envelope of *S. coelicolor*, we chemically isolated so-called sacculi ^37,38^. We then used cryo-ET in combination with a Volta phase plate (VPP) to increase the low frequency contrast of our data^39^.

Tomograms of some of the sacculi displayed extensive folding perpendicular to the long axis of the hyphae. These folds are similar in orientation compared to the shears and tears previously observed in sacculi preps of *Bacillus subtilis* ^40^, likewise supporting a circumferential orientation of the glycan strands in the cell wall (Figure S1)^40^. The cell wall of the purified WT sacculi has a thickness of approximately 30 nm, similar to the width of the cell wall of *B. subtilis* sacculi 40. Furthermore, the tomograms of the *S. coelicolor* sacculi showed that the apex of the tip region appears less densely packed with cell wall material (Figure 1C, D). The subapical parts (Figure 1E) exhibit a higher contrast compared to the apical parts (Figure 1F). Of the nine datasets we have recorded, three tips indeed showed a striking difference in electron density between the apical and subapical regions (Figure 2A). In the other tips the apical regions were not conspicuously different compared to the subapical part. Four other tips were too closely located to the carbon support film and could not be further analyzed. This data set indicates a structural difference between the apex, where the incorporation of new PG into the existing cell wall takes place, and the relatively older cell wall located subapically.

**Figure 2.**
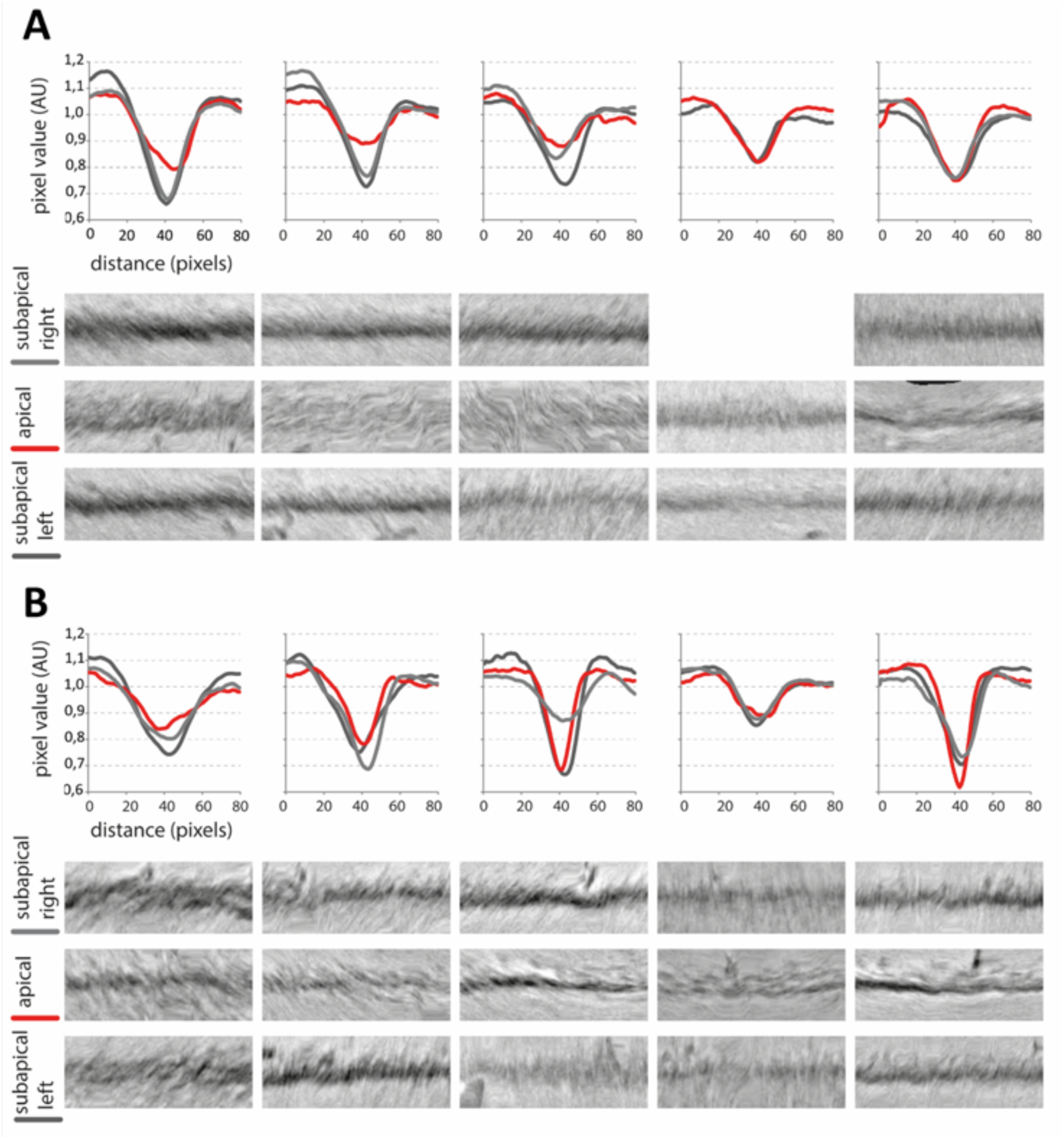
Density profiles of the apical and subapical regions of isolated sacculi. Cryo-electron tomographic slices of five non-treated (A) and five HF-treated (B) *S. coelicolor* sacculi were used to analyze the density of the cell wall. For each sacculus, the apical region (red) together with the subapical regions on the left (dark gray) and right side (light gray) were segmented and straightened. The density was determined by averaging pixel values (arbitrary units) along the straightened cell wall regions. These pixel values are presented in the density plots relative to the distance perpendicular to the cell wall. Representable fractions of the segmented and straightened cell walls are depicted below the density plots.

### Teichoic acids provide structural support to the cell wall architecture of growing tips

To study the contribution of TAs to the overall cell wall architecture, we treated sacculi with hydrofluoric acid (HF). HF cleaves phosphodiester bonds and is a proven method to remove TAs from the cell wall ^9,41,42^. Tomograms of HF-treated sacculi showed no apparent density differences between apical and subapical regions (Figure 2B, n=8). This indicates that the differences in cell wall density between the lateral and apical regions appear to depend, at least in part, on TAs.

The analysis of the HF treated sacculi revealed another difference compared to the non-treated samples: in the absence of TAs, the structure of the sacculi was overall less uniform and revealed a patchy and rough cell wall structure (Figure 3). This frayed appearance of the cell wall was evident in all regions of the sacculus and not restricted to a specific area. In three out of twelve HF-treated sacculi, the sacculus consisted of two distinct lamellae (Figure 4). The other nine HF-treated sacculi did not reveal distinct layers but revealed a more patchy cell wall structure. The patchy cell wall pattern was also evident in a tomogram that we collected from an emerging side branch (Figure S2), suggesting that it correlates to young hyphae. Taken together, these results indicate that the cell wall of *S. coelicolor* is comprised of two distinct lamellae that are structurally linked by teichoic acids.

**Figure 3.**
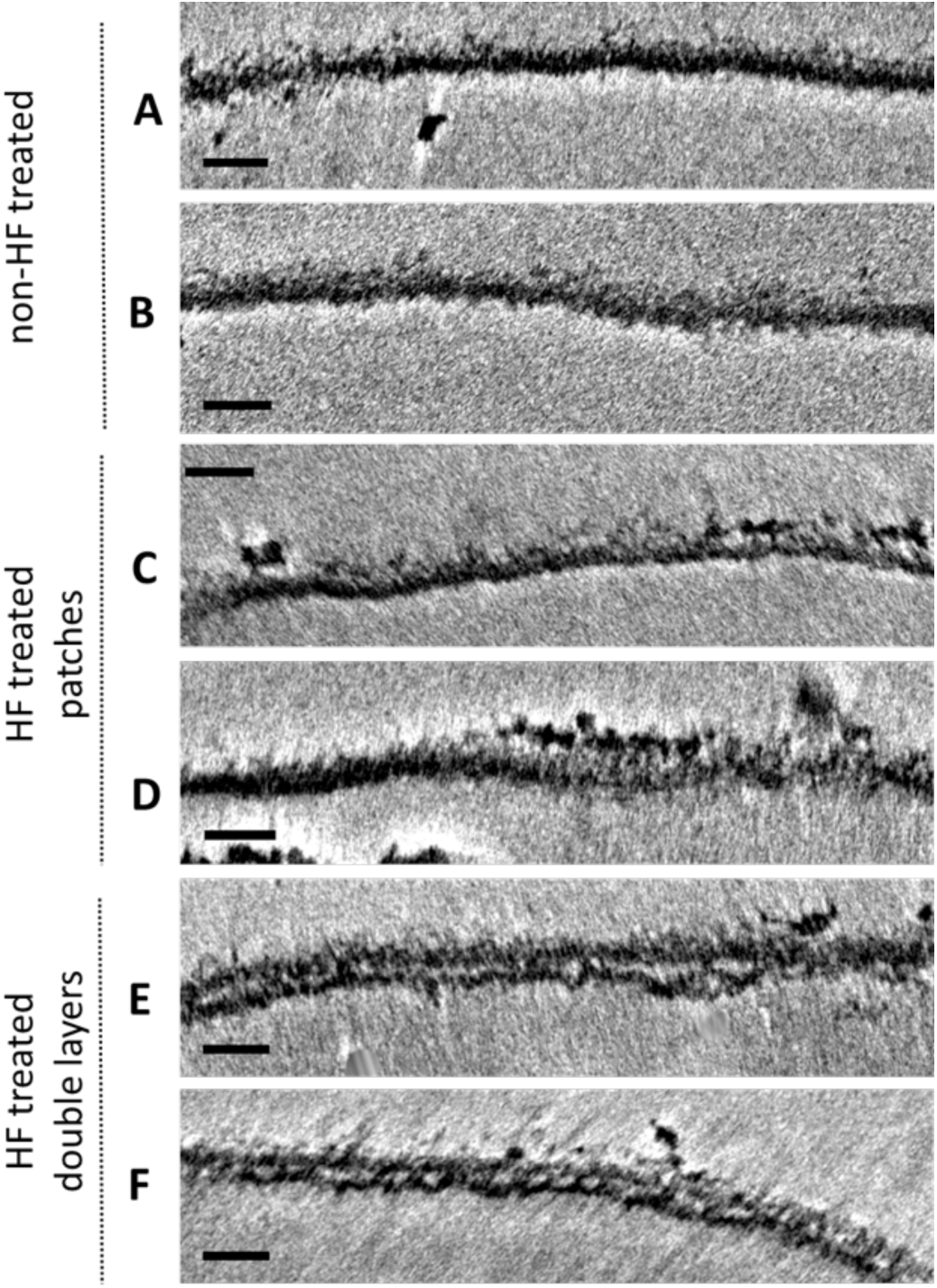
Cryo-ETs of *S. coelicolor* sacculi show alterations in the PG layer upon HF treatment. Panel A & B show a representative part of the PG layer of a non-HF treated *S. coelicolor* sacculus. Panel C-F show representative parts of the PG layers of HF-treated sacculi, classified as patches (C-D) and double layers (E-F). All scale bars are 50 nm.

**Figure 4.**
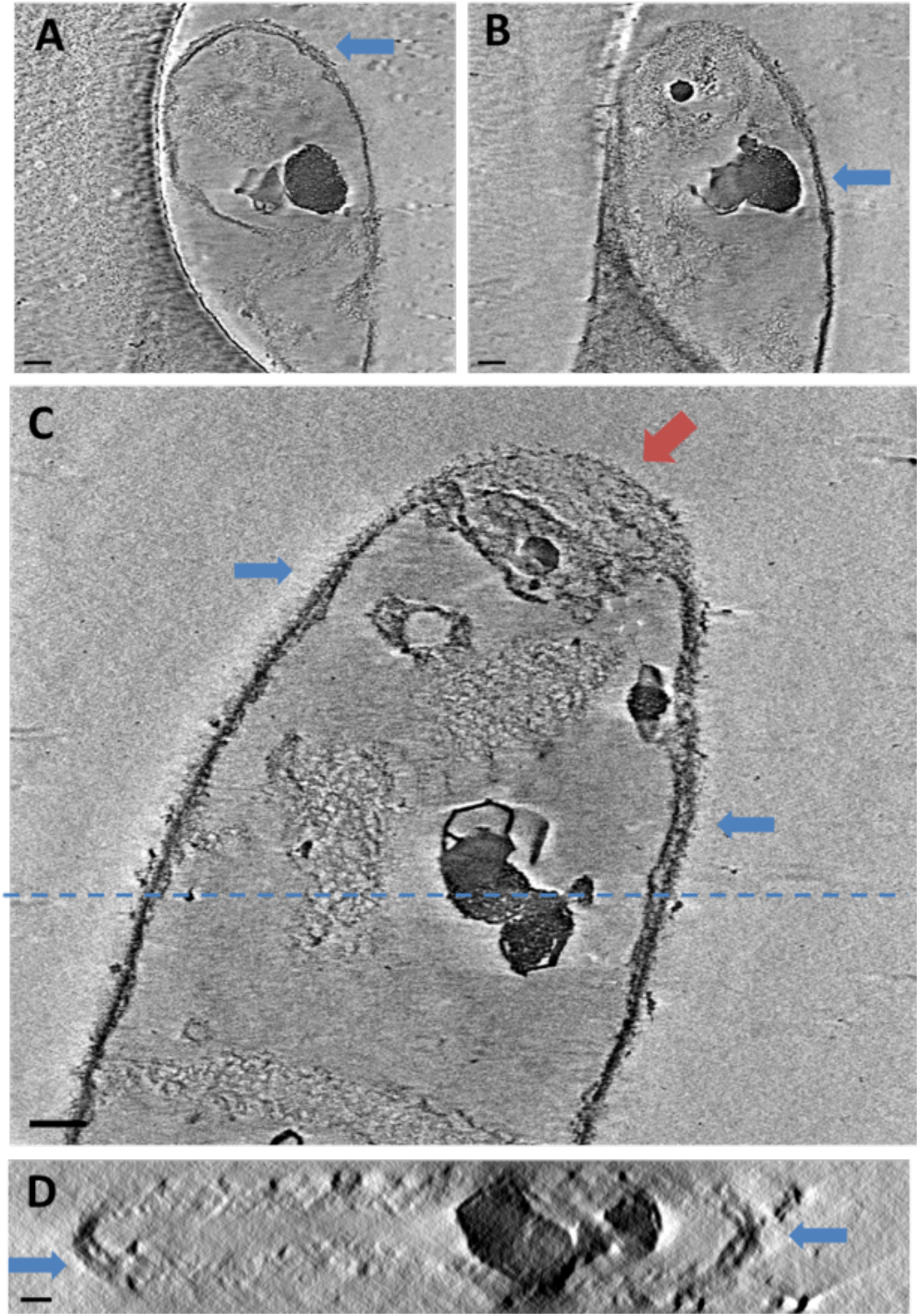
Cryo-ETs of HF-treated *S. coelicolor* sacculi show double layers of PG. The micrographs in panel A & B are the same sacculus, displayed at different Z heights, the blue arrows highlight the presence of a double layer. The micrographs at panel C shows a collapsed double layer at the apex, highlighted by the red arrow. The double layer of the sacculus in panel C can also be seen in a ZX plane (panel D) at the approximate Y height depicted in panel C by the dashed line. Scale bars 100 nm

### Extracellular glycans form a distinct second lamella in the *S. coelicolor* cell wall

To investigate whether the patches and double layers are composed of PG, HF-treated sacculi were treated with mutanolysin that cleaves β-N-acetylmuramyl-(1→4)-N-acetylglucosamine linkages in PG ^43,44^. Exposure to mutanolysin indeed degraded most of the sacculi within 15 minutes and after 90 minutes most individual sacculi were absent (Figure S3). This confirms that the PG is the shape-determining component of the sacculus but does not yet provide an insight into the causes of the patches and distinct layers observed with cryo-ET.

When grown in liquid medium, *S. coelicolor* germlings produce an extracellular matrix leading to self-aggregation into dense pellets.

This pellet structure remains intact after chemical isolation of the sacculi and even after the mutanolysin treatment as observed by light microscopy (Figure S3).

The extracellular matrix is composed of a large variety of glycans, such as poly-N-acetylglucosamine (PNAG) and a cellulose-like polymer, produced by MatAB and by CslA, respectively ^30–32^. The interaction of the cellulose-like glycan and PNAG with the bacterial cell wall on a single cell level has so far not been described. But since the pellet structure is preserved even after sacculi isolation, these polymers might play a direct role in the cell wall architecture of *S. coelicolor*.

To determine the role of these extracellular glycans in the cell wall architecture of *S. coelicolor*, we isolated the sacculi of two mutants strains: *S. coelicolor* M145 Δ*matAB* and Δ*cslA*. Both mutant strains have a clear open morphology compared to the dense cultures grown by the parent *S. coelicolor* strain (Figure S4). The data acquired from the HF-treated *matAB* and *cslA* mutants appeared similar to the sacculi of the WT at first glance (Figure 5 A-D). However, we did not observe patches and double layers in either mutant. Furthermore, comparison of these mutants with the WT revealed a significant difference in cell wall thickness. The WT sacculi had an average thickness of 30.16 nm (± 2.65, n=9). The thickness of the HF-treated WT sacculi showed an average thickness of 31.19 (± 7.12, n=9) and a higher variance, presumably caused by the patches and double layers in this sample group. In contrast, the sacculi of the Δ*matAB* and Δ*cslA* strains showed an average thickness of 23.75 (±5,16, n=9) and 22.78 nm (± 8.29, n=7), respectively. These values significantly differ from the WT (p < 0.01) and the HF-treated WT (p <0.05) when analyzed using a Student’s t test. The remarkably thinner cell wall of both the *matAB* and *cslA* null mutants indicates that the extracellular glycans produced by the MatAB proteins and CslA comprise a considerable part of the cell wall itself. Moreover, the absence of patches and lack of double layers in these mutants imply that these patches are composed to a large degree of these glycans, and that these glycans are thus an integral part of the cell wall.

**Figure 5.**
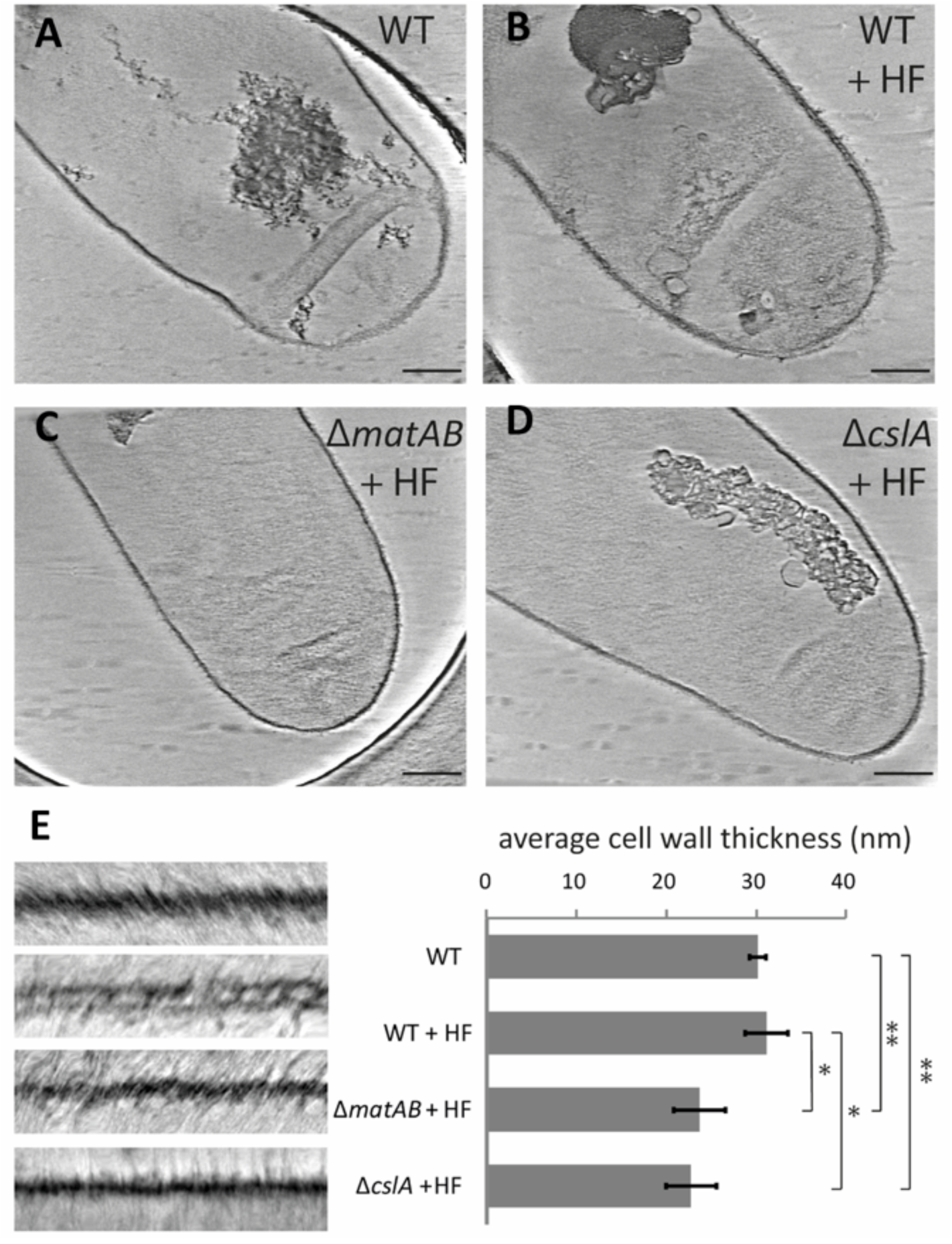
Sacculi of *S. coelicolor* Δ*matAB* and Δ*cslA* appear thinner. The micrographs in panel A-D show a sacculus of *S.coelicolor* WT non-treated (A), WT HF-treated, Δ*matAB* HF-treated (C) and Δ*cslA* HF-treated (D), scale bar 200 nm. The thickness of the overall saccculi are measured and shown in the graph (E), of which representative fractions of the sacculi are depicted in the panels on the left. Graph: * p < 0,05, ** p < 0,01 (Students t-test).

## Discussion

In this work, we show that the cell envelope of the polar growing bacterium *S. coelicolor* is a complex structure composed of PG and extracellular glycans that are structurally linked by teichoic acids Our observations indicate that the TAs directly add to the structural integrity of the *S. coelicolor* cell wall. Chemical removal of the TAs reveals the existence of two distinct lamella, or likely remnants thereof, appearing as a patchy cell wall. These observations were not restricted to the apical region and were also seen subapically. In contrast, the HF-treated sacculli of *cslA* and *matAB* null mutants lacked any observable lamellae, strongly suggesting that lamellae are composed of - or depend on - the extracellular matrix synthesized by the CslA and MatAB proteins. The results presented in this study warrant a revisited model for the cell wall of the polar growing bacterium *S. coelicolor* (Figure 6).

**Figure 6.**
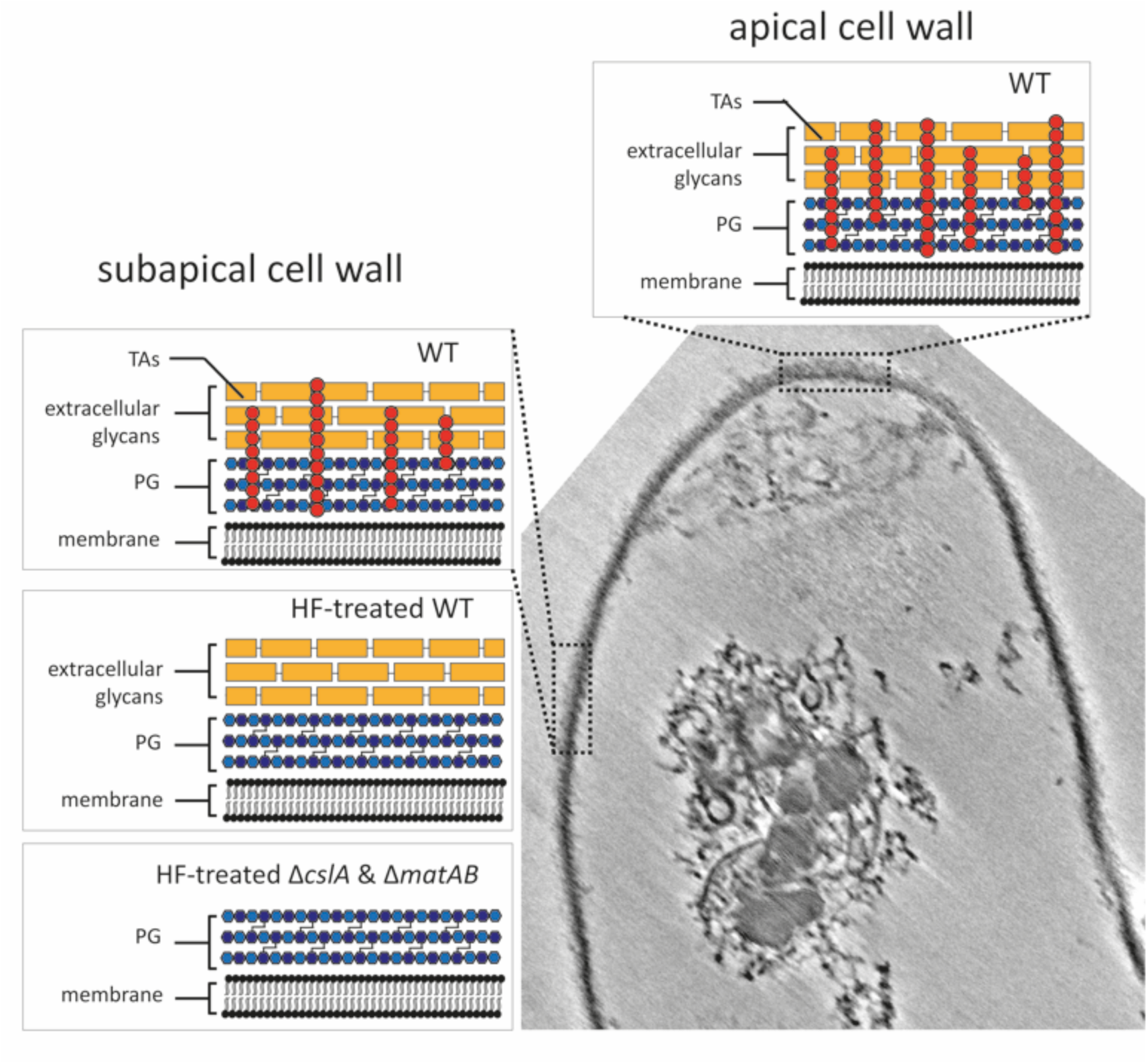
Schematic model of the cell wall architecture of *S. coelicolor*. Micrograph of a WT *S. coelicolor* sacculus, with insets showing the schematic model of the cell wall architecture resulting from the data presented in this study. The model shows the cellular lipid bilayer or membrane, covered by a layer of peptidoglycan (PG) and extracellular glycans, which are interconnected by the wall teichoic acids (TAs). At the top, the apical cell wall panel represents the apex, where new PG is incorporated and cross-linked into the sacculus. The left panels show the subapical cell wall of the WT, HF-treated WT and mutant strains. The HF-treatment causes the removal of the wall teichoic acids, whereas extracellular polymers are absent in the mutant strains Δ*cslA* and Δ*matAB*. The Gram-positive cell wall is composed of the combined structure of wall teichoic acids, extracellular polymers and peptidoglycan.

Cryo-ET has been previously used to study for example cell division events in *Streptomyces* ^45^. However, this is the first study investigating the cell wall structure in this organism. Our results align well with previous cryo-ET work in respect to the orientation of the glycan strands in the sacculus. In these studies, the glycan strand orientation in the PG of the laterally growing diderm bacteria *Escherichia coli* and *Caulobacter crescentus* ^46^ and the monoderm *Bacillus subtilis* ^40^ were shown to be circumferentially positioned around the bacterium. None of these studies reported a difference between the apical and lateral parts of the sacculi. In contrast, we show here that the apical regions appear less-densely packed compared to the lateral regions in *S. coelicolor*. However, this difference was not found in all apical regions, which is likely due to the fact that not all tips are actively growing. Additionally, this difference is also not apparent in the HF-treated sacculi, which supports a growth model where the TAs may play an important role.

Our data further indicate that TAs play an essential role in the structural integrity of the cell wall by tethering the distinct lamellae of the cell wall to one another. The role of the TAs in the cell wall structure has been previously studied in the Gram-positive pathogens *Staphylococcus aureus* and *Listeria monocytogenes* ^47^. Both pathogenic bacteria showed alterations in the PG layer as a result of treatment with the antibiotic tunicamycin, which inhibits the WTA biosynthetic enzyme TagO ^48^. Classical negative stain and sectioning TEM showed that the PG layer of *L. monocytogenes* and *S. aureus* appears thinner and rougher upon tunicamycin administration ^47^. The micrographs from these studies show a striking resemblance with our data of the *S. coelicolor* sacculi treated with HF. Additionally, early cryo-EM research on frozen-hydrated sections of *B. subtilis* cell wall fragments also showed that removal of the TAs lead to a loss of rigidity and reduced thickness by around 10 nm ^49^. This finding correlates well with the loss of integrity we observed upon TAs removal from the *S. coelicolor* cell wall.

The data presented in this study show that the *S. coelicolor* sacculus is thinner in the absence of TAs and the glycans produced by the CslA and MatAB proteins. This could indicate that both glycans form an additional rigid layer providing protection during apical growth. Both glycans are important for pellet-formation in *Streptomyces* and are associated with biofilm formation in other bacteria ^50–52^.

In summary, the results presented here indicate that the cell wall of polar growing bacterium *S. coelicolor* contains the structurally important components PG, TAs and extracellular glycans that together compose a thick and complex cell wall. In our model, we propose that the cell wall is composed of a layer of densely cross-linked PG, with a layer of extracellular glycans produced by *cslA* and *matAB*-encoded proteins on top and exposed to the exterior of the cell. These layers are packed together by wall TAs, which are interweaved throughout the cell wall in hyphal tip. These findings lead to the insight that the *S. coelicolor* cell envelope is a complex network composed of PG and extracellular glycans, and that is structurally interlinked by TAs.

## Methods

### Strains

*Streptomyces coelicolor* A3(2) M145 was obtained from the John Innes Center Strain Collection. The deletion mutants of *cslA* (SCO2836) and *matAB* (SCO2963 and SCO2962) in *Streptomyces coelicolor* M145 used in this study have been previously published by Xu *et al.* 2008 ^53^ and Van Dissel *et al.* 2015, 2018 ^30,31^ respectively. All techniques and media used to culture *Streptomyces* are described in ^54^. Spores of *Streptomyces* were harvested from Soy Flour Mannitol (SFM) agar plates. Fresh spores were used to inoculate 400 mL of Tryptic Soy Broth supplemented with 10% (w/v) sucrose, in 2 liter flasks with coiled coils. The liquid cultures were grown while shaking at 200 rpm and 30°C for 12 hours prior to sacculus isolation.

### Sacculus isolation

Sacculi of *S. coelicolor* WT, Δ*matAB* and Δ*cslA* were isolated using a protocol based on the method of B. Glauner ^55^. Liquid cultures of 12 hours old were resuspended in cold TrisHCl pH 7.0, subsequently boiled in 4% sodium dodecyl sulfate (SDS) for 30 min and washed with milliQ. The sample was enzymatically treated with DNase, RNase and trypsin. Then the sample was again boiled for 30 min in 4% SDS and washed. The sample was pelleted and resuspended in 48% hydrofluoric acid (HF), which is shown to be sufficient to quantitatively remove teichoic acids from the sacculus ^56^. After 48 hours of HF treatment, the sample was washed multiple rounds and concentrated.

### Cryo-electron tomography

Prior to vitrification of the sample, 10 nm colloidal gold beads (Protein A coated, CMC Utrecht) were added to the sacculi suspension as fiducial markers in a 1:20 or 1:50 ratio. Vitrification was performed using a Leica EM GP plunge freezer. The EM-grids were glow-discharged 200 mesh copper grids with an extra thick R2/2 carbon film (Quantifoil Micro Tools). 3,5 µl of the sample was applied to the EM grid and blotted at a temperature of 16°C and a humidity between 95-99%. Grids were automatically blotted with a blot time of 1 second and plunged into liquid ethane.

Data was collected using a Titan Krios instrument (Thermo Fischer Scientific) equipped with a 300 keV electron gun, Volta phase plate and Gatan energy filter with K2 Summit DED (Gatan, Pleasanton, CA). The sample was tilted −60, + 60, and imaged with 2 degrees increment, cumulative exposure of 100 electrons and a pixel size of 4.2848 Å. Tilt series were acquired using Tomography 4.0 software (Thermo Fisher Scientific) with usage of the Volta phase plate (Thermo Fisher Scientific) and a defocus set to −0.5 µm. The tomograms were reconstructed by applying a weighted back-projection algorithm with SIRT-like filtering, using IMOD software ^57^.

### Mutanolysin treatment and microscopy

HF-treated sacculi of *S. coelicolor* were kept in mutanolysin buffer containing mutanolysin. The sample was incubated at 30 degrees Celsius, and at three time points (0, 15 and 30 min upon start of the treatment) sample was taken for observation with a light microscope and prepared for room temperature TEM. For bright field microscopy, 3 µl of the sample was placed on a glass microscopy slide with cover slip and observed with a Zeiss Axio Lab A1 upright microscope with an Axiocam MRc camera. For TEM, approximately 20 µl of the sample was applied on a 200 mesh copper with continuous carbon grid and left to dry at room temperature. The sample was observed using a 120 kV Talos TEM (FEI/Thermo Fisher) equipped with Lab6 filament and Cita CCD camera.

### Data analysis

The density plots and thickness measurements of the cryo-ET data was performed with the open-source software ImageJ and (FIJI) plugins ^58^. Per sacculus, the approximate middle was set by determining the slice where the sacculus is the broadest. Approximately 50-100 slices around the sacculus was summed to one micrograph, as the sacculi can contain wrinkles or does not always lay flat in one plane. From the summed micrograph, the sacculus was traced using various filters and skeletonization of the micrograph using tools in FIJI. The skeletonized line following the sacculus was used as selection to straighten the sacculus, which was subsequently used for a vertical density/profile plot using the Profile Plot tool in FIJI. For the tip vs. lateral region comparison, a selection of 800 pixels (approximately 679 nm) from the most curved region was designated as the tip region. The density plot of this region was compared to the density plots of adjacent (left and right, if available) lateral regions. The cell wall thickness per sacculus was determined by subtracting the background signal from the density plot values of the straightened cell wall. The average background pixel value was determined by averaging all pixel values of the regions outside and inside the sacculus selection.

## Acknowledgements

We thank Julio O. Ortiz (Netherlands Center for Electron Nanoscopy (NeCEN), Leiden, the Netherlands) for support on cryo-ET data acquisition with the Volta Phase Plate and critical reading of the manuscript. And we thank J. Willemse (Institute of Biology, Leiden University, Leiden, the Netherlands) for his help with FIJI for data analysis. We thank Prof. Justin Nodwell (University of Toronto) for critically reading the manuscript.

This work has been supported by iNEXT, PID:2265, funded by the Horizon2020 programme of the European Commission.

## SUPPLEMENTARY FIGURES

**Figure S1.**
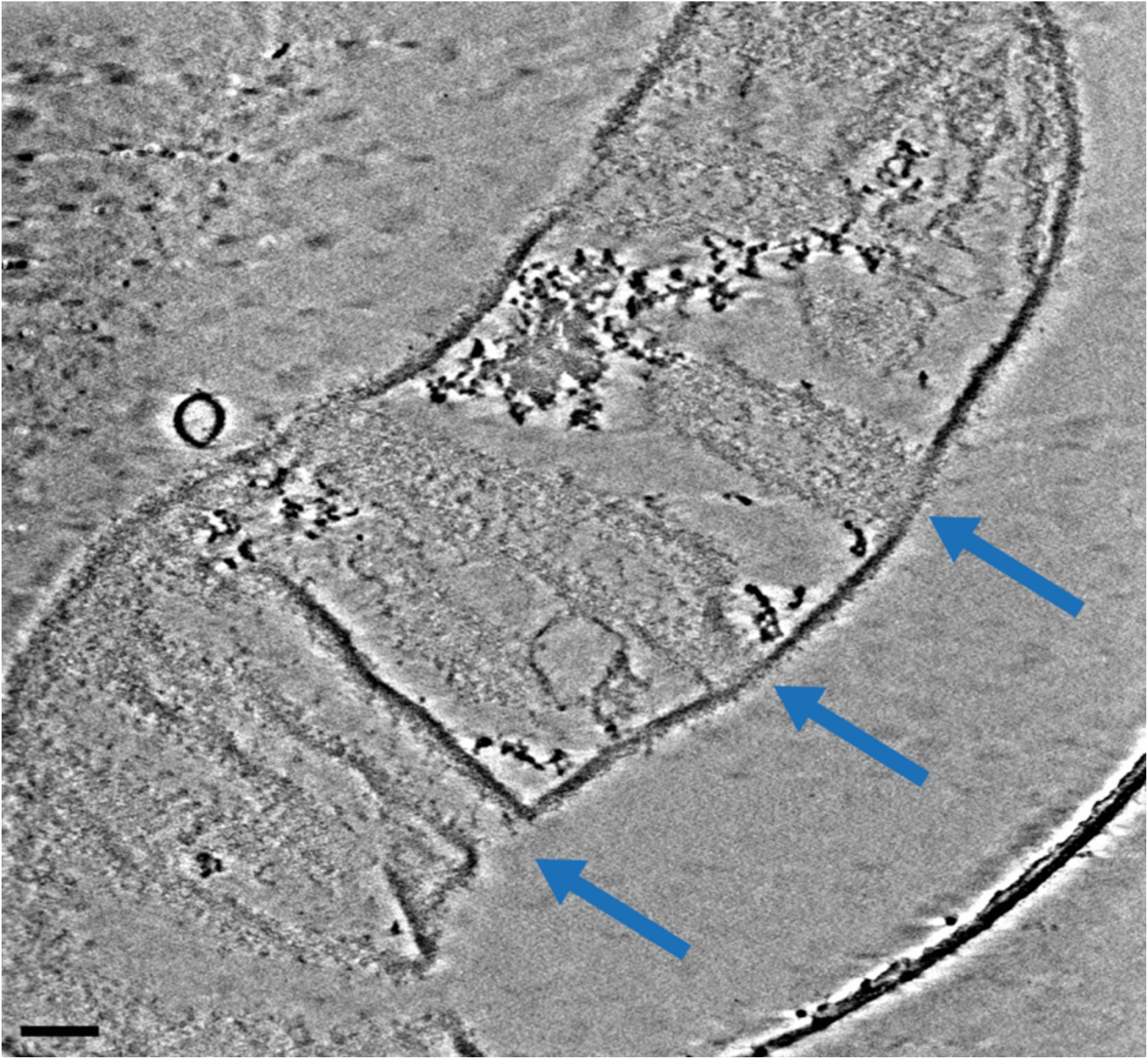
Cryo-electron tomograms of isolated *S. coelicolor* sacculus. Example of a cryo-electron tomogram of an *S. coelicolor* sacculus showing multiple folds perpendicular to the cellular axis. Folds are highlighted by the blue arrows. Scale bar 100 nm.

**Figure S2.**
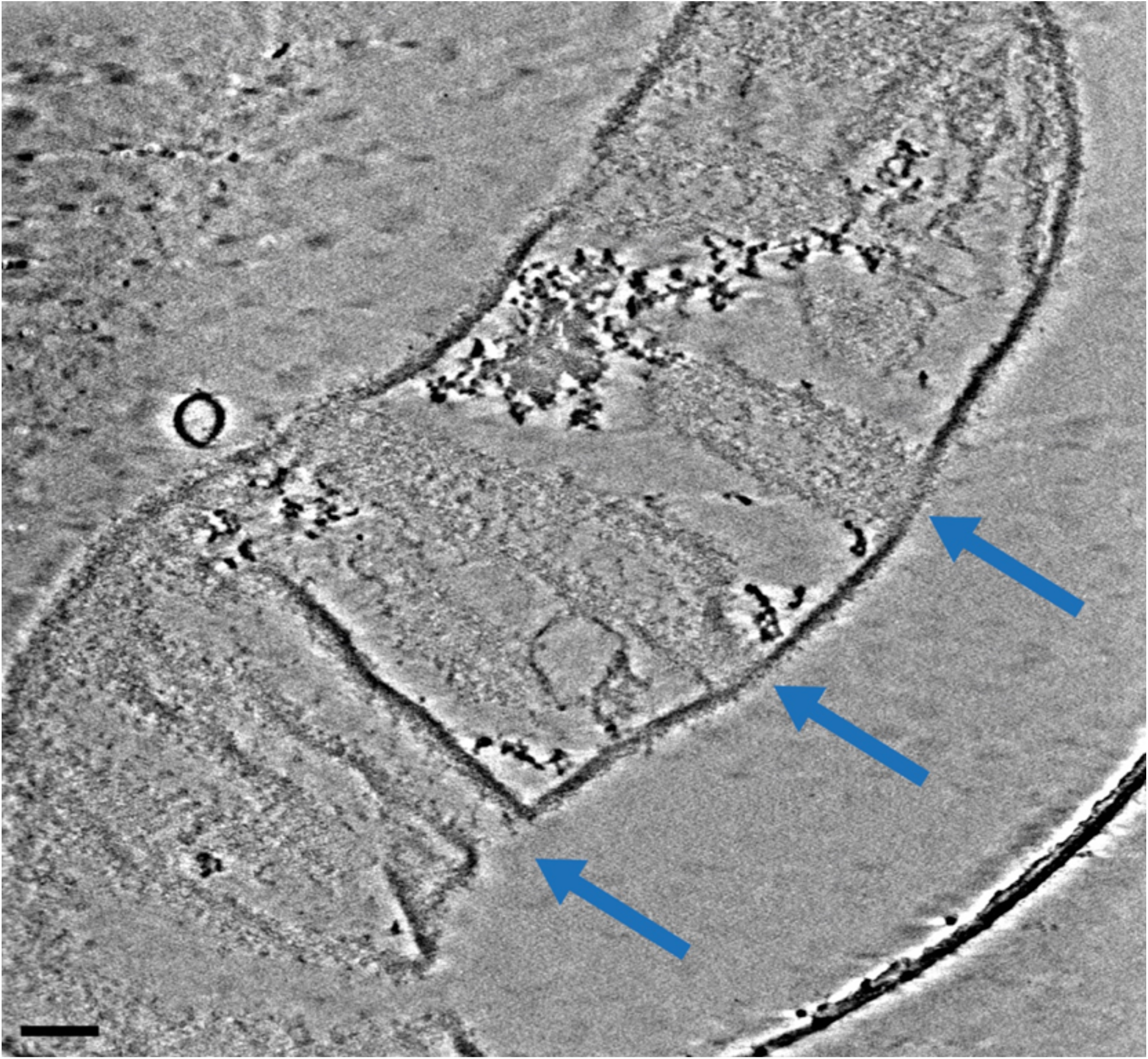
Cryo-ETs of HF-treated *S. coelicolor* sacculus with a side-branch. Cryo-ET micrograph of a side-branch. The two large panels are micrographs taken from two different Z heights (A, B) to accurately compare the cell wall morphology of the folded sacculus. The insets on the right side are from the lateral sides of the hyphae (bottom and top smaller panels) and of the tip and lateral side of the newly formed branch (middle panels). Scale bars 100 nm, insets 20 nm.

**Figure S3.**
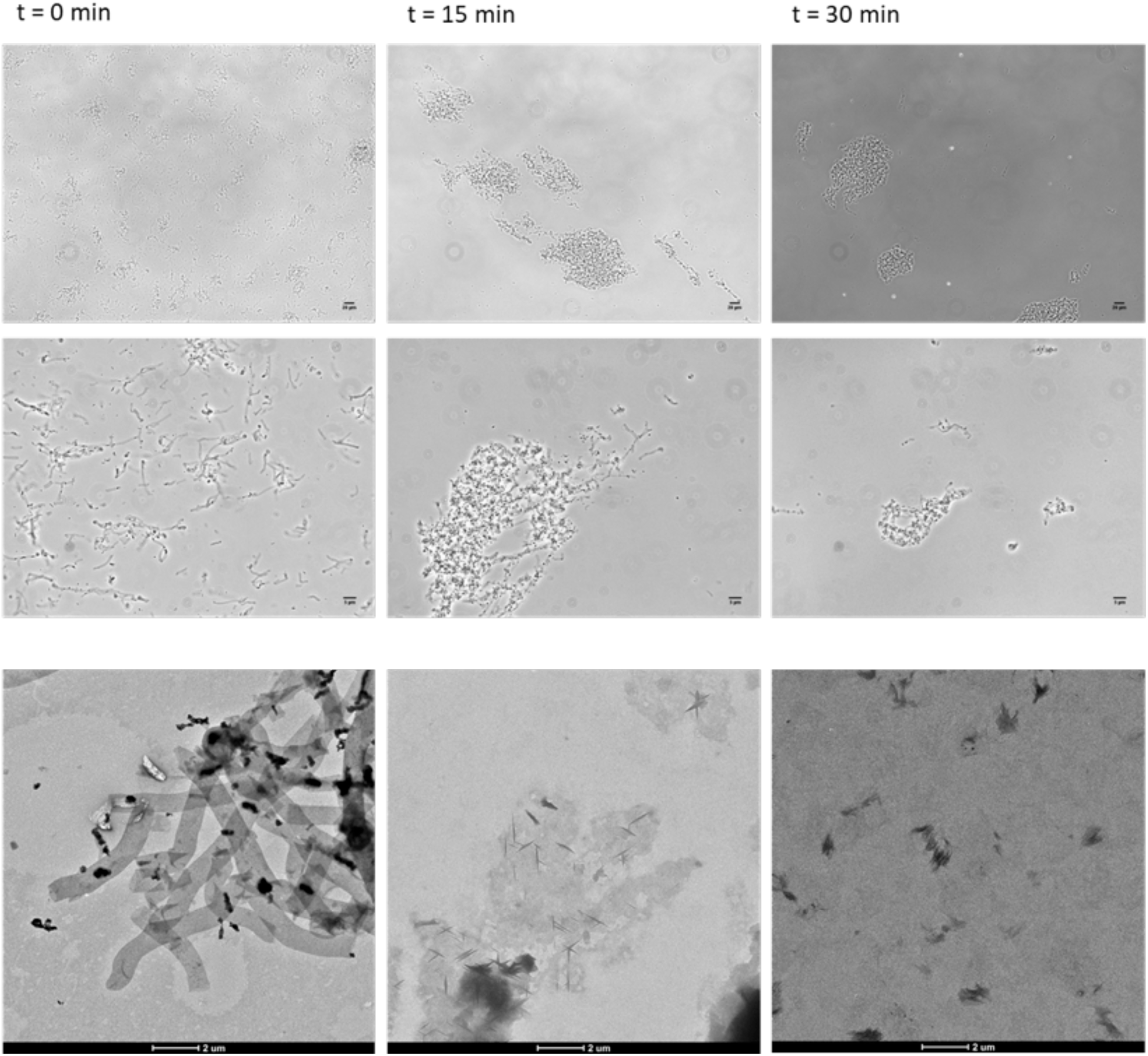
HF-treated *S. coelicolor* sacculi degrade upon mutanolysin treatment. The HF-treated *S. coelicolor* sacculi were incubated with mutanolysin for 0, 15 and 30 minutes. The upper two rows show light microscopy micrographs, whereas the bottom row shows room temperature EM to visualize the degradation of the sacculi upon mutanolysin treatment.

**Figure S4.**
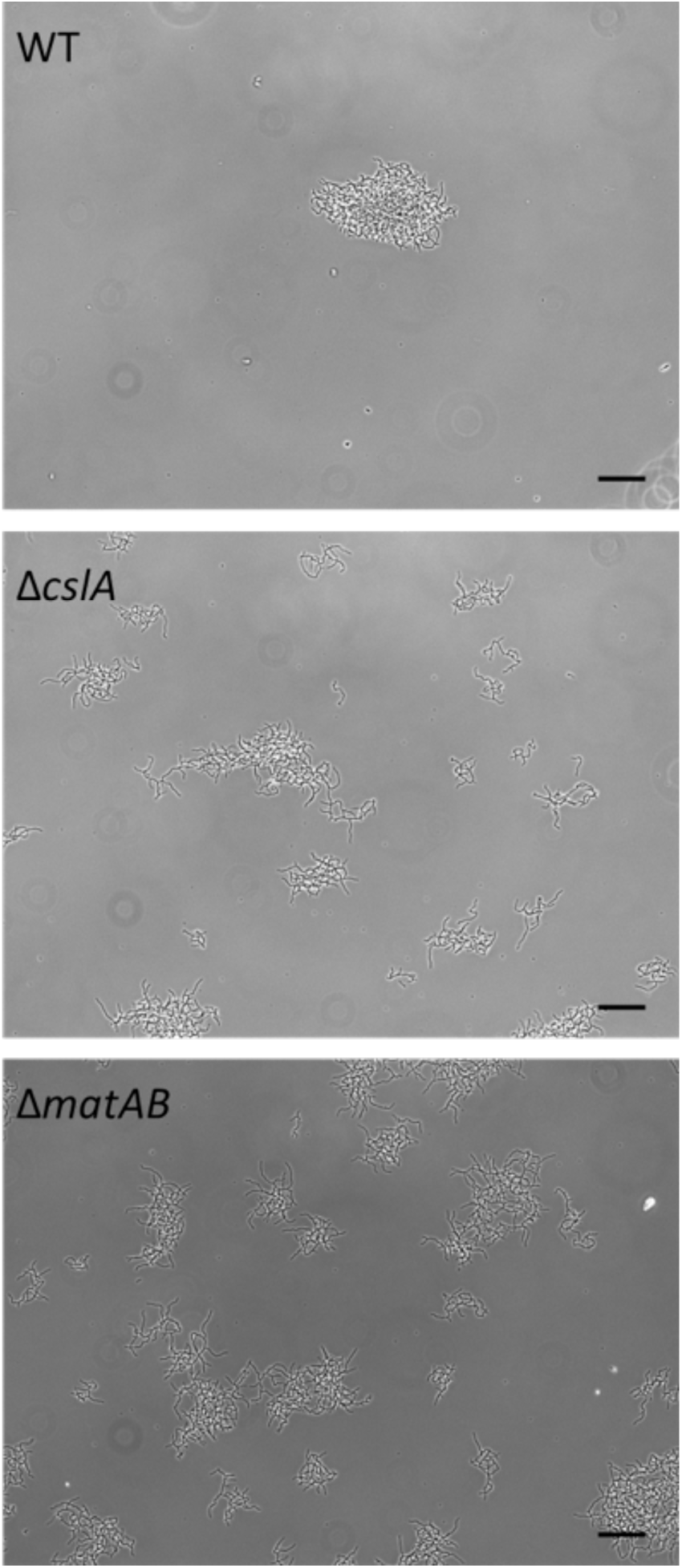
Pellet morphology of *S. coelicolor* WT, Δ*cslA* and Δ*matAB*. Pellet morphology of *S. coelicolor* WT, Δ*cslA* and Δ*matAB* after 12 hours of growth in liquid TSBS medium. The WT forms a dense pellet, whereas Δ*cslA* and Δ*matAB* show a dispersed morphology. Scale bars 50 µm.

